# Ecology of low pathogenicity avian influenza virus H7 in wild birds in south-eastern Australia prior to emergence of high pathogenicity avian influenza H7 in poultry

**DOI:** 10.64898/2025.12.08.693062

**Authors:** Michelle Wille, Paul Eden, Silvia Ban de Gouvea Pedroso, Ian G. Barr, Allison Crawley, Kyeelee Driver, Victoria Jimenez, Peter D. Kirkland, Peter T. Mee, Matthew J. Neave, Kim O’Riley, Andrew J. Read, Vittoria Stevens, Teri Visentin, Andrew C. Breed, Marcel Klaassen

**Affiliations:** WHO Collaborating Centre for Reference and Research on Influenza, at the Peter Doherty Institute for Infection and Immunity; Centre for Pathogen Genomics, Department of Microbiology and Immunology, University of Melbourne, at the Peter Doherty Institute for Infection and Immunity.; Wildlife Health Australia, Canberra, Australian Capital Territory, Australia; Department of Primary Industries and Regions, South Australia, 5000, Australia; City and Environment Directorate, Canberra, Australia Capital Territory 2601, Australia; Elizabeth Macarthur Agriculture Institute, NSW Department of Primary Industries and Regional Development, Menangle, New South Wales 2568, Australia; Agriculture Victoria Research, AgriBio Centre for AgriBioscience, Bundoora, VIC 3083, Australia; Commonwealth Scientific and Industrial Research Organisation (CSIRO), Australian Centre for Disease Preparedness, Geelong, Victoria 3320, Australia; Centre for Integrative Ecology, School of Life and Environmental Sciences, Deakin University, Geelong, Victoria 3217, Australia; Department of Agriculture, Fisheries and Forestry, Canberra, Australia Capital Territory 2601, Australia

## Abstract

Adding to the global burden of high pathogenicity avian influenza (HPAI) H5N1, an unprecedented five HPAI H7 outbreaks occurred globally in 2024. Of these, three occurred in southeast Australia, with the independent emergence of HPAI H7N9, H7N8, and H7N3, resulting in the destruction of 2 million poultry. Historical data demonstrates that H7 outbreaks in Australia do not occur randomly, rather, there is a strong association between the timing of the previous H7 outbreaks and rainfall patterns in southeastern Australia. We aimed to address a hypothesis wherein prior to H7 outbreaks in poultry, there was a detectable change in H7 prevalence and/or virus diversity in wild bird populations. We addressed this using virological and serological surveillance data generated from multiple programs. Despite the collection of thousands of samples, there was only weak evidence to support our hypothesis, which provides strong incentive to evaluate current surveillance approaches for the purposes of risk prediction. However, in alignment with a previous analysis, there is strong support for a relationship between H7 outbreak probability and rainfall patterns across southeast Australia. Overall, improved understanding of the ecology and evolution of H5 and H7 viruses in wild bird reservoirs is pivotal to global disease preparedness and response.

## Introduction

Wild birds, and waterfowl in particular, are the main reservoirs for nearly all influenza A viruses [1]. The vast majority of avian influenza viruses (AIVs) are low pathogenicity (LPAI) and wild birds have no or limited clinical signs of disease when infected, a likely result of long-term co-evolution. With the advent of high-density poultry production, high pathogenicity avian influenza (HPAI) viruses, have increasingly emerged [2,3]. These HPAI viruses may develop in poultry after LPAI H5 and H7 viruses spill over from wild birds into poultry [2]. HPAI viruses pose a major threat to poultry health, and more recently, wild birds and mammals, livestock and humans [4]. While HPAI H5 viruses, notably those caused by goose/Guandong lineage HPAI H5N1 are currently in the limelight, HPAI H7 viruses continue to emerge. Since 1963, 38 HPAI H7 outbreaks have occurred globally [2], causing substantial costs to poultry industries. For instance, >33 million poultry were depopulated due to HPAI H7 outbreaks from 2015-2020 [5]. Beyond poultry, in China, H7N9 was identified as a substantial public health threat, causing >1500 human infections between 2013-2019 [6].

While HPAI H5 viruses remain absent from Australia, twelve HPAI H7 outbreaks have occurred in Australian poultry since 1976 [7–10]. These outbreaks have not been evenly spaced over time, with 2024 a remarkable outlier: three of the five H7 outbreaks globally occurred in southeastern Australia [9,11–13]. These three outbreaks in Australia occurred from May to July 2024, with a HPAI H7N9 impacting one property near Terang, Victoria, a HPAI H7N3 infecting seven properties in the Golden Plains Shire, Victoria, and a HPAI H7N8 affecting six properties in the Hawkesbury area of New South Wales and spreading to two properties in the Australian Capital Territory [9]. In 2025, yet another independent HPAI H7N8 outbreak occurred near Euroa, Victoria [10]. There is a convincing correlation between the timing of historic HPAI and LPAI H7 outbreaks in poultry and rainfall patterns in southeastern Australia [8,14]. This correlation is thought to be caused by a cascading effect of rainfall on waterfowl numbers and movements, LPAI epidemiology, and spill-over risk to poultry. Increasing amounts of water in the landscape during prolonged periods of rain lead to improved breeding conditions for waterfowl in ephemeral wetlands, driving a considerable increase in immunologically naïve juvenile birds. Inevitable droughts follow these rainfall events, leading to the disappearance of ephemeral wetlands and waterfowl migrating to and aggregating in permanent water bodies, which are predominantly situated in more cultivated regions of the country also including dams on poultry farms. These conditions facilitate an increase in AIV prevalence in wild waterfowl, alongside a potential increased likelihood of AIV spillover to poultry [8,14,15]. While the correlation between rainfall patterns and AIV outbreaks in poultry is convincing, the predictive power of the relationship is limited. For instance, not *all* periods of increased rainfall followed by periods of drought have led to outbreaks in poultry. Thus, there is demand for improved clarity into the factors contributing to heightened likelihood of spillovers of AIV from wildlife into poultry, benefitting risk mitigation. One way of attaining this could be by augmenting the rainfall-based predictions by integrating information on prevalence and subtype diversity in wild birds. Specifically, we hypothesize that in the period prior to these outbreaks there may be both increased rainfall as well as an increased prevalence and diversity of LPAI H7Nx in the wild waterfowl populations.

To address this hypothesis and attempt to better understand factors in the wild bird reservoir preceding spillover and emergence of HPAI H7 outbreaks in poultry, we used viral prevalence and seroprevalence data collected in ducks in southeastern Australia over the period 2022-2024. Specifically, we integrated data from multiple sources, included both molecular surveillance and serology, which were investigated for both overall AIV and H7-specific prevalence, and analysed viral sequence data. Furthermore, we revisited the role of rainfall, as a predictor of H7 outbreaks in poultry. Together, the compilation of these data provided a weak signal in surveillance data that suggested an increased potential risk of a LPAI H7 spill-over from wild birds into poultry in the period leading up to the outbreaks of 2024. While there may be a clear benefit in using a combination of environmental data (i.e. variations in water in the landscape and, therewith, the availability of duck breeding habitat; cf Ferenci et al. 2021) and wild bird AIV monitoring to predict HPAI outbreak risk and support risk mitigation, this can only be achieved by modifying structured risk-based surveillance in order to detect fine scale patterns in virus dynamics.

## Methods

### Sample collection and sample screening from live captured wild ducks in Victoria

Samples were collected from live, apparently healthy wild ducks, caught between June 2022 and July 2024. Wild ducks were caught using baited funnel walk-in traps and cannon nets. Baited funnel walk-in traps were deployed on land or in very shallow water allowing for foraging by dabbling ducks. Traps baited with wheat and/or corn were operated between dawn and dusk. Cannon nets to capture roosting ducks were operated during day. During winter, wild ducks were captured in four regions in Victoria: i.e. at the Western Treatment Plant of Melbourne Water (144.57°E, -38.00S), and in the areas around the cities of Geelong (Serendip: 144.41°E, -38.00°S; Wallington: 144.53°E, -38.23°S), Wangaratta (Milawa: 146.45°E, -36.49°S; Tarrawingee: 146.54°E, -36.35°S; Thoona: 146.11°E, - 36.36°S) and Sale (Heart Morass :147.2°E, -38.14°S). In summer, ducks were only captured in Wallington.

Individual samples from live caught wild ducks were collected as previously described [16]. Briefly, both oropharyngeal and cloacal swab samples were collected using a sterile tipped applicator and placed together into a vial with virus transport media (VTM, brain heart infusion [BHI] broth-based medium [Oxoid] with 0.3 mg/ml penicillin, 5 mg/ml streptomycin, 0.1 mg/ml gentamicin, and 2.5 g/ml, amphotericin B). Following collection, swab samples were kept cool (4-8°C) for up to 4 days prior to being stored at -80°C. Up to 200µl of blood was collected from the brachial vein using the Microvette capillary system for serum collection (Sarstedt). Following blood-sample collection, samples were kept cool (4-8°C) for 7-14 hours, followed by centrifugation to separate sera according to manufactures recommendations.

Samples collected were screened as previously described [16]. Briefly, in 2022 and 2023, RNA was extracted using the NucleoMag Vet Kit (Scientifix) on the Kingfisher Flex. Extracted RNA was subsequently assayed for a short fragment of the matrix gene [17] using the SensiFAST Probe Lo-Rox qPCR Kit (Bioline). During winter 2024 samples were assayed by the Victorian Department of Economic Development, Biosciences Research Division. RNA was extracted using the MagMax 96 Viral Isolation Kit (Applied Biosystems, Thermo Fisher Scientific) using the Kingfisher Flex (Thermo Fisher Scientific). The RNA was tested using the AIV Type A real-time hydrolysis probe (TaqMan®) based RT-qPCR assay, which targets the highly conserved matrix (M) gene [18]. RT-qPCR reactions were performed using the AgPath-ID^TM^ One-Step RT-PCR kit (Thermo Fisher Scientific). Positive samples were confirmed and subsequently assayed for H5/H7 using a qPCR approach at Agriculture Victoria (2022, Ct<35) and/or the Australian Centre for Disease Preparedness (2023-2024, all positives) using accredited methods.

### Detection of anti-NP antibodies in live captured wild ducks

Serum samples were assayed using the Multi Screen Avian Influenza Virus Antibody Test Kit (IDEXX, Hoofddorp, The Netherlands) following manufacturer’s instructions. An S/N value <0.6 was considered positive, as per [19,20]. Next, serum samples positive by anti-NP ELISA which had sufficient remaining sample volume were subjected to a hemagglutination inhibition (HI) assay using 1% chicken red blood cells as previously described [20]. Briefly, sera were treated with*Vibrio cholerae* receptor-destroying enzyme (RDE II; Denka Seiken Co., https://denka-seiken.com), then inactivated with 1.5% sodium citrate. For the haemagglutination inhibition assays (HI), treated sera was added with a starting dilution of 1:10. Antiserum against A/Emu/VIC/20-03243-3/2020(H7N6) was used as positive control. The HI was run with both A/Emu/VIC/20-03243-3/2020 (H7N6) and A/grey teal/Victoria/18985/2024(H7N6) as antigens.

### National Avian Influenza Wild Bird (NAIWB) molecular surveillance data

To complement the serological and molecular results from our live captured duck program, we requested data from the (Australian) National Avian Influenza Wild Bird (NAIWB) surveillance program [21,22]. We included data from the south-eastern states and territories in which the Murray Darling Basin is situated because historically the HPAI H7 outbreaks have predominantly occurred in or in close proximity to the Murray Darling Basin [2,9]. However, not all sampling sites in each state is within the Murray Darling Basin. States and territories include: Victoria, South Australia, New South Wales and Australian Capital Territory. We requested national data in alignment with the sampling time frame of our study: 2021-2024. We requested two types of data: (1) qPCR results and (2) H7-HA sequence data. For the qPCR results, we requested the number of samples collected, number of matrix positives, and number of H7 qPCR positives at each sampling event. The vast majority of samples are from environmental faecal samples, targeting wild ducks. However, for most samples there is limited detail and/or certainty on the wild bird species sampled. NAIWB data also included samples collected from hunter-shot ducks. For the sequence data, we requested the metadata required for generation of virus designations, as well as date of collection. Sample collection, testing and sequencing approaches used by the NAIWB partners are described in [22,23]. The requested data was received in September 2024.

NAIWB test results included data collected on both individual and pooled swabs (up to three samples per pool). For the calculation of prevalence data, where a pooled test result was positive, it was assumed that only one of the three pooled swabs caused this positive result.

### Statistical analysis

For the serology analysis of the live captured ducks, we considered seroprevalence (i.e. number of anti-NP antibody positive samples divided by total number of samples taken), and H7 seroprevalence (i.e. number of samples with H7 antibodies as a fraction of total number of samples taken). We used a similar approach for AIV and H7 prevalence data (qPCR) of live captured ducks. These four different sets of prevalence data were analysed using generalized linear mixed effect models with a binomial response using function “glmer” within the “lme4” package. In the models we used species and a combination of year and season as fixed factors, while we controlled for potential regional differences by using region as a random factor. Only data for the species of which overall sample size equalled or exceeded 10 individuals were considered in the analysis. For season, we divided the data into spring-summer (Oct-Mar) and autumn-winter (April-Sept). Year in which the season started was also included in this fixed factor. The significance of the fixed factors was evaluated using Type II anova (i.e. the effect of a factor was tested after correcting for the effects of all other factors in the model), using function “Anova” within the “car” package. Where a factor had a significant effect on prevalence, pairwise posthoc multiple comparisons were conducted for that factor using the emmeans package.

Analysis of the AIV and H7 prevalence in the NAIWB qPCR data was similarly approached, however, while year and season were similarly defined as for the live-captured duck data set, we used year and season as two separate factors rather than one combined fixed factor. We used state in which samples were collected and analysed (Victoria, New South Wales and South Australia) as random factor, where samples from the Australian Capital territory (ACT) were included with the New South Wales data as the ACT is geographically nested within New South Wales. Bird species was not included as an explanatory variable in the analysis as that information was not available with certainty for the environmental faecal swabs collected. Finally, we also analysed live-captured ducks and NAIWB data sets together using the same approach as for the NAIWB data in isolation.

Data analysis was performed in R version 4.4.1 integrated into R Studio version 2024.04.2. Graphs were made using the ggplot2 package within R.

### Virus isolation of samples from live captured ducks

Positive samples were inoculated into 11-day old embryonated chicken eggs via the allantoic route. Following incubation for 3 days, allantoic fluid was harvested and tested for agglutinating activity using an haemagglutination assay (HA) assay. RNA was extracted using the Qiagen QiaAMP Virus Kit, and the subtype confirmed through sequencing.

### Full genome sequencing and phylogenetic analysis

Full genome sequencing was undertaken as previously described [23]. Briefly, samples were sequenced on an Illumina MiSeq with up to 24 samples pooled per sequencing run by use of dual-index library preparation and the Nextera XT DNA Library Preparation kit and 300-cycle MiSeq Reagent v2 kit (Illumina). Sequence reads were trimmed for quality and mapped to respective reference sequence for each influenza A virus gene segment using Geneious Prime software (www.geneious.com) (Biomatters, Auckland, NZ)

We incorporated all full length H7 sequences generated in Australia since 2015, including those published in Wille *et al.* 2022 (2015-2020), those requested from the NAIWB (2021-present), and the genome generated in this study. Sequences were aligned using MAFFT [24] integrated within Geneious Prime. We performed a linear regression of root-to-tip distances against year of sampling with TempEst [25]. Subsequently, a time-scaled phylogenetic tree was estimated using BEAST v1.10.4 [26], under the uncorrelated lognormal relaxed clock [27] and SRD06 codon structured nucleotide substitution model [28], and the Bayesian skyline coalescent tree prior [29]. One hundred million generations were performed, and convergence was assessed using Tracer v1.8 (http://tree.bio.ed.ac.uk/software/tracer/). A maximum credibility lineage tree was generated using TreeAnnotator following the removal of 10% burn-in, and were visualised using Fig Tree v1.4 (http://tree.bio.ed.ac.uk/software/figtree/). Effective population size was estimated using the “Skygrid” workflow (https://github.com/Wytamma/skygrid), with a six-year grid and 12 transition points per year

For other segments, two maximum likelihood trees were constructed. First, “global” trees using backbones reported in [30] to demonstrate phylogenetic placement relative to globally circulating viruses. Second, “local” trees comprising only the top 10 blast hits, retrieved in Geneious Prime (17 January 2024), as well as a North American outgroup sequence retrieved from the BV-BRC database (https://www.bv-brc.org/). In both cases, maximum likelihood trees incorporating the best-fit model of nucleotide substitution were estimated using IQ-Tree [31], and 1000 ultrafast bootstraps. Trees were visualised using Fig Tree v1.4

### Rainfall and outbreaks

We updated the analysis conducted by Ferenczi et al. (2021) using additional rainfall and AIV outbreak data in poultry until January 2025. Monthly total rainfall across the Murray Darling Basin (mm/month) between January 1970 and January 2025 were downloaded from the Bureau of Meteorology, using http://www.bom.gov.au/cgi-bin/climate/change/timeseries.cgi?graph=rain&area=mdb&season=allmonths&ave_yr=0.

For details on the analysis see Ferenczi *et al*. (2021). In brief, we used a generalised linear model to conduct a logistic regression across all months. An outbreak month was scored as a 1 and a non-outbreak month as 0 as dependent variable. To appropriately weight co-occurring outbreaks (i.e. outbreaks happening within the same months of the same year), one of the outbreaks was moved to the following month. We ran 600 models where in each model we used rainfall as the explanatory variable calculated for a different combination of rainfall period (varying from 1-24 months) and time-lag period (varying from 0-24 months). For each model of the total of 600 (=24 rainfall periods x 25 time-lag periods) models the AIC was calculated. The top performing models were classified as the models where the AIC was within two AIC units of the model with the minimal AIC.

## Results

### Virus prevalence in wild ducks of southeast Australia

To reveal the dynamics of H7 in waterfowl of southeastern Australia, we leveraged two datasets. First, we live caught and collected samples from 402, 148 and 347 wild ducks in the autumn-winter (here defined as April – September) of 2022, 2023, and 2024 respectively. In all years, most samples were collected in June and July (n=962) such that in 2024, the majority of samples (n=335) were collected between 12 June and 26 July 2024 (*i.e*. after the first HPAI H7 outbreak in poultry in 2024, Fig S1). Thirty-four samples were collected from live caught wild ducks in the summer half year (October - March) of 2023-2024. Across all years, samples were collected from as many as 10 different species (or hybrids), including Grey teal (*Anas gracilis*), Chestnut teal (*Anas castanea*), Pacific black duck (*Anas superciliosa*), Australian shelduck (*Tadorna tadornoides*), Wood duck (*Chenonetta jubata*), Australasian shoveler (*Anas rhynchotis*), Hardhead (*Aythya australis*), Pink-eared duck (*Malacorhynchus membranaceus*), Pacific black duck x domestic duck hybrid, and Chestnut teal x Pacific black duck hybrid (Data available at (https://github.com/michellewille2/H7Manuscript). Data from the NAIWB program comprised both hunter-shot ducks (n=1063), and faecal environmental samples (n=9024) (Table S1).

AIV (matrix qPCR) prevalence in live-caught birds ranged from 0 – 12.5% across the seasons (Fig 1A, Fig S2; see figures for 95% confidence intervals of all prevalence estimates). In the statistical analyses of these data, only samples from Grey teal, Chestnut teal, Pacific black duck and Australian shelduck were considered, as all other species/hybrids had only small sample sizes of n<8. AIV prevalence varied significantly across the four season/year periods (Х^2^(3)=16.1, P<0.002), with AIV prevalence in autumn-winter 2022 being significantly higher than in autumn-winter 2024 (P<0.001). In both 2022 and 2023, AIV prevalence was ∼12%. In 2024 autumn-winter, AIV prevalence was only 4%. No samples were positive of the 34 collected in the spring-summer. There was no significant effect of species (Х^2^(3)=2.5, P=0.467).

**Figure 1:**
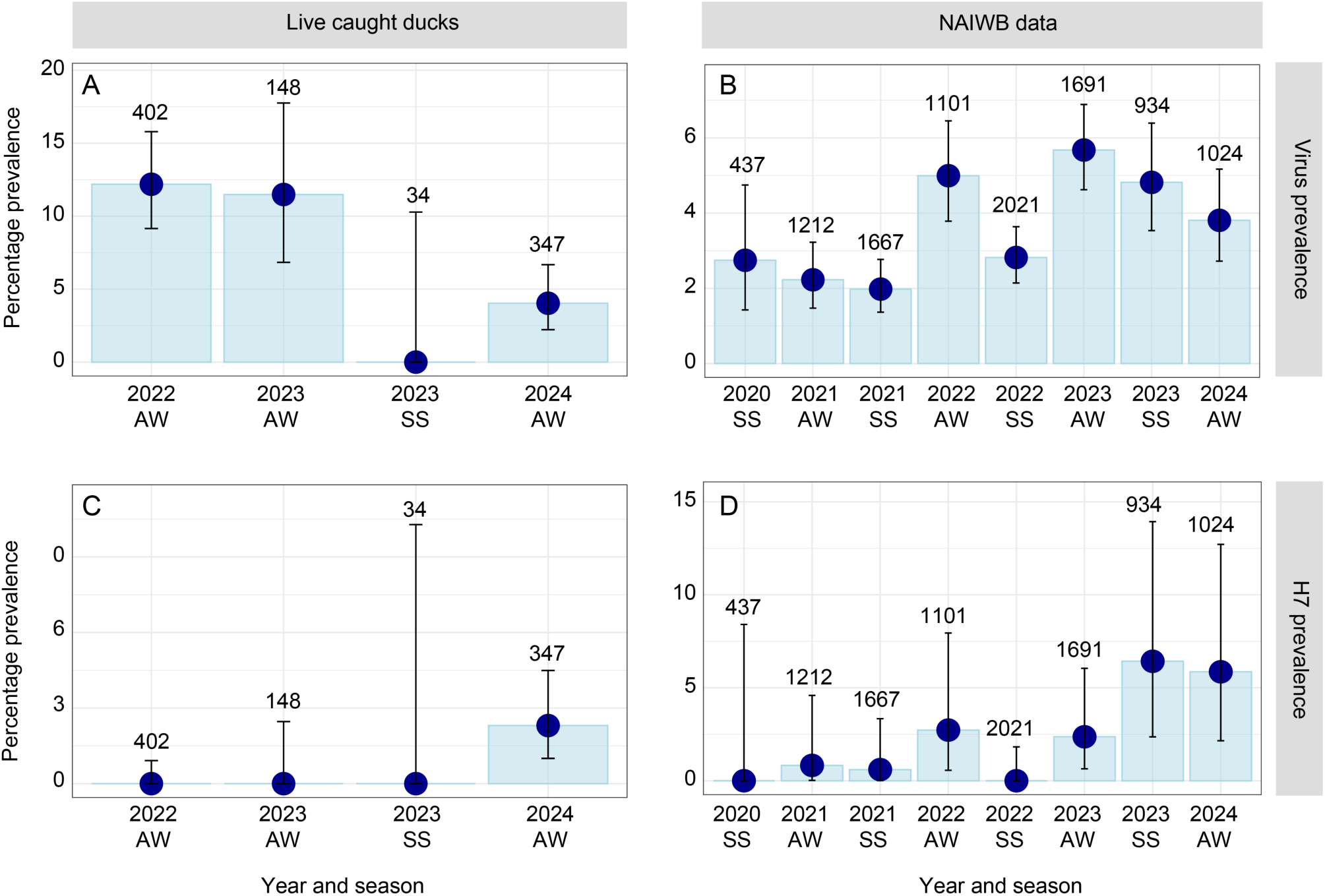
Seasonal and annual variations in AIV prevalence in live caught wild ducks in Victoria and from NAIWB data based on environmental and hunter-shot birds. (A) AIV prevalence, based on matrix qPCR assays in live caught birds in Victoria (B) AIV prevalence in samples collected from the NAIWB program. NSW: New South Wales (which includes data from Australian Capital Territory), SA: South Australia, Vic: Victoria. (C) H7 prevalence in live caught birds, where positives are those identified based on H7 qPCR assays, and “negative” comprise all samples assayed for avian influenza matrix qPCR. (D) H7 prevalence in the NAIWB data. Bars are 95% confidence intervals, values are number of samples. AW (autumn-winter) runs from April-September, and SS (spring summer) from October-March.

AIV prevalence in the NAIWB data was 4% on average (Fig 1B). Season (autumn-winter versus spring summer) had a significant effect on AIV prevalence (Х^2^(1)=7.4, P<0.007), with autumn-winter prevalences being on average higher than spring-summer prevalences. Also, year had a significant effect on AIV prevalence (Х^2^(4)=20.9, P<0.001), with 2021 being significantly lower than 2022 and 2023 and, 2024 being significantly lower than 2023. Combining all data (i.e. for live caught birds and NAIWB) provides a similar picture that does not suggest that AIV prevalence was substantially higher in the lead up to the 2024 outbreaks.

H7 prevalence was substantially lower than AIV prevalence. In live-caught ducks in Victoria, no H7 viruses were found across all seasons and years investigated, except in the period where the outbreaks took place (2024, autumn-winter), when it was nearly 2.5% (Fig. 1C). However, there were no statistically significant differences across seasons/years and species across the live caught birds. The H7 prevalence across the much larger NAIWB data set was <1% across the investigated seasons and years (Fig. 1D). Season had no effect on H7 prevalence (Х^2^(1)=0.02, P=0.879), but year did (Х^2^(4)=9.6, P<0.047), with post-hoc testing providing near-significant effects between the years 2021 and 2024 (P=0.09). Combining the data from NAIWB and the live caught birds makes the year effect in H7 prevalence substantially stronger (Х^2^(4)=20.9, P<0.001) with significant differences between 2022 and 2024 (P<0.006) and 2023 and 2024 (P<0.02), and near-significant differences between 2021 and 2024 (P=0.066). Overall, the differences across years are small and we did not observe an increase in H7 prevalence prior to the outbreaks.

### Viral diversity in wild ducks of south-eastern Australia

A substantial diversity of LPAI viruses were detected in wild ducks. Eighteen HA-NA subtype combinations were detected through sequencing from live captured ducks in 2022 and 2023: H1N3, H2N3, H2N7, H2N9, H3N6, H3N8, H3N9, H4N1, H4N6, H4N8, H5N3, H6N1, H6N9, H8N5, H10N3, H10N7, H11N3, H11N6. Analyses of H4, H5, and H10 viruses have been reported previously [30,32]. Further analysis of these data are outside the scope of this study. Critically, in neither 2022 nor 2023 did we recover H7 genomes.

We successfully isolated one H7 virus from live caught ducks in 2024, and the full genome was successfully sequenced from both the original sample (presented here) and isolate A/grey teal/Victoria/18985/2024(H7N6). We were not successful in isolating or sequencing other H7 positive samples, likely due to high Ct values (Ct 34-39). The HA sequence of A/grey teal/Victoria/18985/2024(H7N6) was 99% similar to both A/chicken/Victoria/10-17/2024(H7N9) (EPI3413127) and A/chicken/Victoria/24-01759-3/2024(H7N3) (EPI3413135) (Fig. 2). The PB2, PB1, PA, NP, NA, and NS (B allele) sequences fall into clades which comprise viruses from Australian ducks. As most recent viruses genomes in GenBank (since 2021) from Australia comprise H4, H5, H10 viruses [30,32], these sequences dominate the most closely related viruses in trees and blast searches. The M segment is mostly closely related to an Australian sequence A/Radjah shelduck/Northern Territory/20231282-03/2023(H5N1)( PP922946), which likely comprises a more recent introduction into Australia as the other top blast hits are of Eurasian sequences (Fig. S3). Two other positive samples from live captured ducks in 2024 were subtyped as H1N2 through blast of recovered genomic fragments, but we were not able to recover complete, high-quality genomes for detailed analysis.

**Figure 2.**
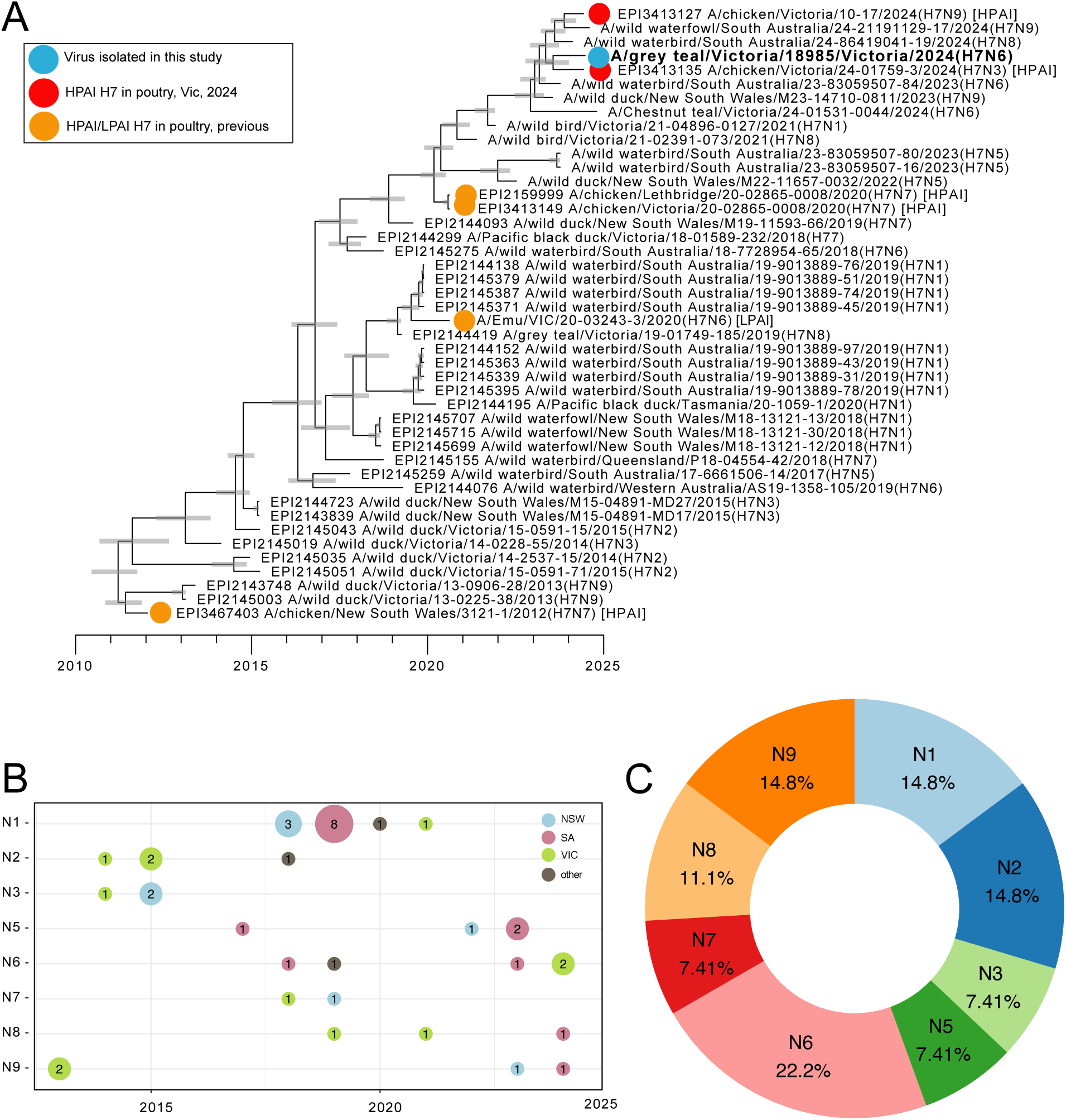
(A) Time structured phylogenetic tree comprising H7 sequences from Australia in public repositories (up till October 2024), the NAIWB program, and generated in this study. Scale bar in years. Bars comprise 95% highest posterior density around node heights. The genome recovered in this study is in blue, HPAI sequences in public repositories are in red (2024) or orange (previous outbreaks). Analysis of other segments of A/grey teal/Victoria/18985/2024(H7N6) is presented in Fig. S3. (B) NA subtypes for all H7 viruses sequenced by the NAIWB program since 2013. Different colours correspond to different states. Bubble size and numbers within the bubbles corresponds to the number of sequences. (C) NA subtypes for all H7 viruses sequenced by the NAIWB program since 2013. Different colours represent different NA subtypes, and the proportion is included within the coloured slice.

Since 2021, 13 H7 genomes have been sequenced from New South Wales, Victoria, and South Australia by the NAIWB program, and all are LPAI (Fig. 2). These H7-HA sequences are within 96% similarity to each other, and all H7-HA sequences from 2024 (n=4) are within 99% similarity. These LPAI H7-HA sequences from 2024 are 99.2% similar to HPAI sequences A/chicken/Victoria/10-17/2024(H7N9) (EPI3413127) and A/chicken/Victoria/24-01759-3/2024(H7N3) (EPI3413135) (which are the two index sequences for the Victorian HPAI outbreaks [9]). To attempt to understand whether there was a larger effective population size of H7 in the waterfowl reservoir prior to the 2020 and 2024 outbreaks, we undertook a skygrid analysis. However, the skygrid precision estimate (smoothing factor) is very high, and the model favours a smooth demographic history, that is, no fluctuations or increases at certain time points. A substantial caveat to this analysis is that we may not have a suitable number of sequences per year to adequately reveal subtle increases and decreases in population size, as evidenced by large 95% Highest Posterior Densities (HPDs) around the estimate (Fig. S4).

In 2024, H7 genomes comprised three different NA subtypes: H7N6 was detected in Victoria (n=2), and H7N8 (n=1) and H7N9 (n=1) were detected in South Australia. Since 2012, 8/9 NA types have been found associated with H7, although there was some heterogeneity in detection between year, state, and sampling event (Fig S5). Overall, if only one sequence per sampling event is incorporated, we find a similar proportion of different NA subtypes, ranging from 7.4% (N3, N5, N7) to 22.2%(N6). N1, N2, N9 occurred in 14.8% of sequences (Fig S5). Overall, there is not a substantial bias towards specific NA subtypes in wild birds, which may explain the diversity of NA subtypes reported in the HPAI outbreaks in 2024.

### Serology: a window into the past

As the detection window for active infection is small (*i.e*. 4-10 days), we investigated changing seroprevalence across our live caught duck cohort. While not different across species (Х^2^(3)=2.1, P=0.556), seroprevalence was significantly different across seasons (Х^2^(3)=129.9, P<0.001) with seroprevalence in autumn-winter 2024 standing out as significantly higher compared to all other investigated seasons (Fig. 3A); from autumn-winter 2022 till spring-summer 2024 seroprevalence was ∼22%, with a sudden increase in seroprevalence in autumn-winter 2024 to ∼60% (Fig. 3A). These data support our hypothesis that the AIV viral burden in ducks was higher in 2024 relative to the two previous years. However, anti-NP-ELISA represents the prevalence of AIV across all subtypes in the population and is not specific to any one specific subtype. Considering seroprevalence against only H7, rather than all AIV, did not appear to have a significant effect on statistical difference across seasons (Х^2^(3)=2.4, P=0.497; Fig. 3B). Yet, species did have a significant effect (Х^2^(3)=12.4, P<0.01), with Australian shelduck having a significantly higher H7 seroprevalence than the two teal species, but not Pacific black duck.

**Figure 3.**
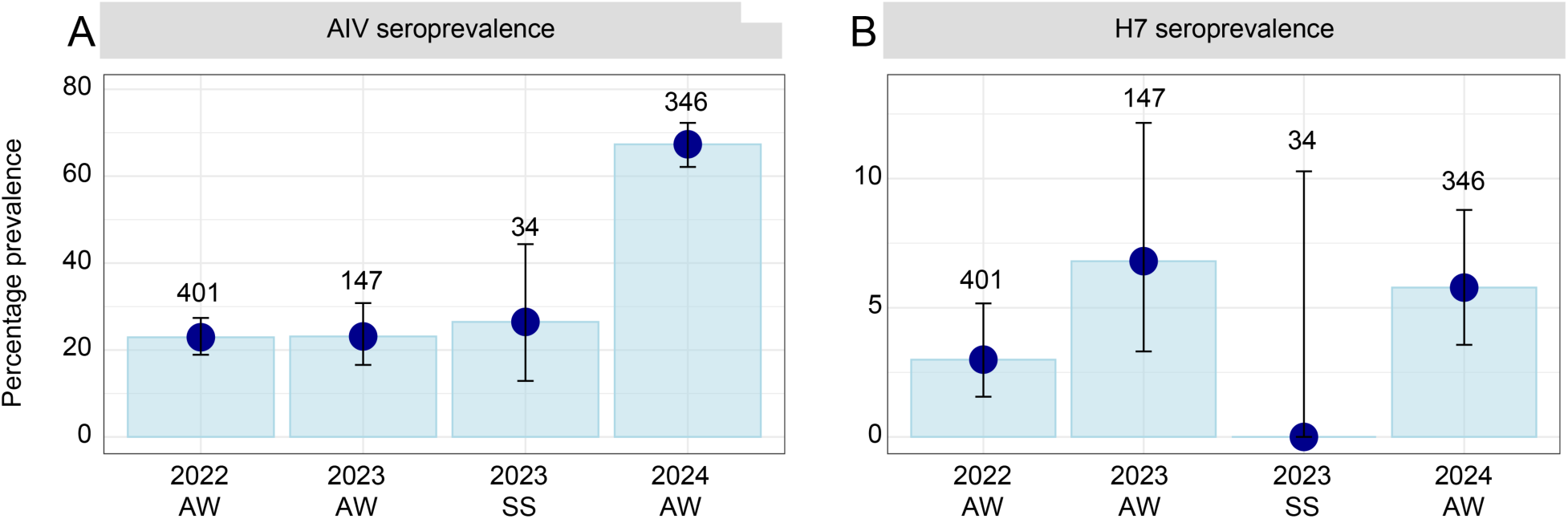
Avian influenza seroprevalence in live captured ducks. (A) Avian influenza seroprevalence, based on anti-NP ELISA, (B) H7 seroprevalence based on HI assays. Prevalence is reported as a percentage. Estimate shown in filled circle. Bars are 95% confidence intervals, values are number of samples. AW (autumn-winter) runs from April-September, and SS (spring summer) from October-March. Corresponding virus prevalence for these data can be found in Fig. 1A, C

### Revisiting correlation between rainfall and occurrence of outbreaks

Ferenczi et al. (2021) demonstrated a link between rainfall and the timing of HPAI and LPAI outbreaks in poultry. Updating their analysis with recent rainfall data across the Murray Darling Basin and AIV outbreaks in poultry in south-eastern Australia, confirms the pattern that periods of high rainfall followed by a period of drought are associated with an increased risk for AIV outbreaks in poultry (Fig. 4). The outbreaks in 2024 were preceded by a substantial increase in rainfall starting in 2022, followed by a profound drying period. In alignment with Ferenczi et al. (2021), our updated re-analysis identified the time lag between the onset of a rainfall period and the maximum chance of outbreaks was a relatively narrow period with a mean of 27.5 months (range 27-28 months) across the top 14 models that were within 2 AIC units. This overall time lag across these top models was composed out of a rainfall period with a mean of 10.8 months (range 7-15 months) and a time lag after rainfall (i.e. the period of drying out) with a mean of 16.8 months (range 13-20 months).

**Figure 4.**
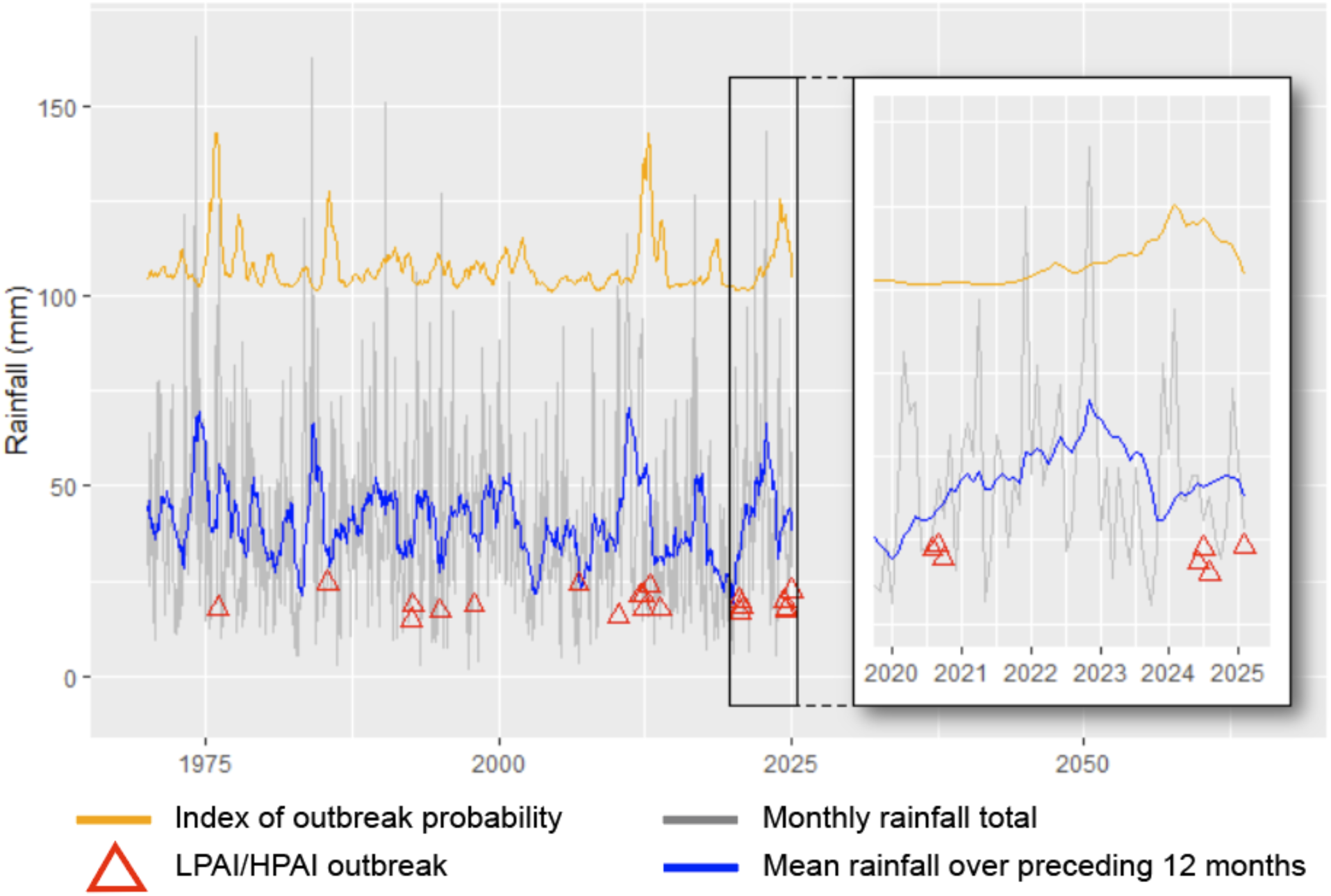
Rainfall in the Murray Darling Basin, HPAI and LPAI outbreaks in poultry (note their prevalent occurrence in periods of drought), and an index of outbreak probability based on the average predicted probability of the top models. Inset zooms in on the period 2020-2025.

## Discussion

Globally, the timing and predictability of HPAI H7 emergence events remains unknown. In 2024-2025, while the world was focussed on HPAI H5, seven HPAI H7 outbreaks occurred [9–11,13,33,34] of which four in Australia. Despite 20 H7 outbreaks in poultry on the Australian continent since 1976 [2], we still do not fully understand the epidemiological factors that drive spillover and, consequently, the emergence of HPAI from LPAI H7 strains. Better identification of the risk factors leading to outbreaks has the potential to be a critical tool for the poultry industry, potentially averting the destruction of millions of birds at a substantial financial cost, including the cascading effects on chicken meat and egg production.

Herein we hypothesized that HPAI outbreaks occurred following a period of increased H7 prevalence in wild waterfowl which, in Australia, is linked to rainfall patterns [8,14]. Despite bringing together a multitude of data types (molecular surveillance, serology, genomic data) from two surveillance programs, we did not detect a clear, easily identifiable increase of LPAI H7 prevalence and diversity in samples collected from wild birds in the Murray Darling Basin preceding the 2024 outbreaks. Rather, all the data from wild birds we gathered and analysed provided only weak evidence for an increased potential of a LPAI H7 spillover from wild birds into poultry leading up to the outbreaks of 2024. It is important to consider, however, that neither surveillance system is designed to detect nuanced (and potentially short-lived) changes in H7 prevalence. Our surveillance in wild ducks occurred for just a few weeks in the winter, and while NAIWB sampling occurs year-round, samples are collected at monthly or quarterly intervals and only for the purpose of virus detection. Our updated analysis on the correlation between H7 outbreaks in poultry and rainfall confirmed that rainfall continues to be the most important predictor of HPAI H7 outbreaks in Australia [8] given the current surveillance approach. Specifically, we again found that the risk of avian influenza (both LPAI and HPAI) outbreaks in poultry increases after a period of high rainfall followed by a period of drought, with the peak risk being approximately two years after the onset of the high-rainfall period. The critical assumptions in this cascade of events is that high rainfall leads to more water in the landscape, increased breeding activity amongst ducks, with a recruitment of many immunologically naïve and susceptible juvenile birds into the population [35]. During subsequent periods with less rainfall, ephemeral wetlands dry out, leading to concentrations of ducks into more permanent wetlands, which lead to higher contact rates and thus higher LPAI prevalence in wild birds, which in turn can increase the risk of spillover into poultry [8]. That we did detect signal from rainfall data, but not from the current approach to wild bird surveillance, highlights the opportunity to strengthen wild bird surveillance to design a fit-for-purpose, systematic integrated surveillance program across both time and space that is capable of detecting fine-scale changes in virus prevalence in wild bird populations. While such systems are labour intensive, they do exist for LPAI, and have been critical in revealing our understanding of LPAI in wild waterfowl in the northern hemisphere [36,37].

Whilst this study identified a signal from rainfall data though not from wild bird surveillance, there is still valuable information that can be obtained from wild bird surveillance. The structure of surveillance systems for avian influenza vary, but the objectives generally comprise one or more of the following broad categories: 1) early warning of increased HPAI risk; 2) evidence of freedom from infection; 3) detection and monitoring of LPAI diversity and evolution to inform risk and management; and 4) increasing understanding of ecology, epidemiology and risk factors [38]. Provision of early warning through detecting changes in spillover risk from the wild bird reservoir, such as a rapid increase in H7 prevalence and diversity in the reservoir hosts in the months preceding spillover events, would require wild bird surveillance to be more adaptive to the unique climatic influences that drive avian influenza dynamics in Australia. This may include sampling at more regular and frequent intervals, over many years (*i.e.* longitudinal surveillance) but particularly following periods of above average rainfall, as well as strengthening harmonisation of sampling approaches. This would provide finer scale data to understand changes in dynamics, such as accuracy of subtype prevalence estimates and variation over time. The benefits of such high frequency sampling is evidenced from studies undertaken in Sweden which allowed for the timely detection of virus arrival into the system, its amplification in the population, as well as its extirpation (*e.g.* [36,39]). However, such approach is labour intensive and costly, such that an evaluation of appropriate intervals should be undertaken in association with the purpose of the surveillance [38]. From a spatial perspective, consideration should be made of movement patterns for wild bird populations. In the temperate northern hemisphere, where waterfowl follow regular migratory routes and time schedules, a surveillance program can be designed to collect samples at sites which are repeatedly visited by waterfowl, whilst ensuring sites are distributed effectively to detect geographical variation. In contrast, waterfowl in Australia are largely nomadic, following water in the landscape, such that there is a lower degree of consistency in patters of where birds are in space and time [40,41]. Finally, target sample sizes may vary depending on sampling purpose. For instance, samples sizes must be sufficiently large to estimate prevalence accurately, be confident that AIV positives are detected for generation of sequence data, or to prove absence of specific AIV types at a desired level of certainty [38]. In all these instances, increasing demands on accuracy are associated with larger sample sizes. In Australia, LPAI prevalence doesn’t occur in consistent cycles [16,35] as it does in the temperate northern hemisphere [*e.g.* [36]. Prevalence may therefore be <2% [22] or higher than 10% (reported here) at any point in the year, requiring large sample sizes at all time points in the year to have confidence that low prevalence subtypes are being adequately detected.

Most avian influenza surveillance systems rely on detecting active infection, but where our active surveillance systems lack the resolution required to make detailed inferences, we garnered substantial insights from the addition of a serology component in our study. As the window for detecting antibodies is much larger (on the scale of months) compared to active infection (on the scale of days to weeks), it provides a window into the past, giving a comprehensive view into AIV at the population level. Serology can play a key role in contextualising the results from active surveillance programs, particularly for subtypes infrequently detected. For example, serology was critical in clarifying the first H14 detection in North America, and paired with active infection results, revealed the spread of this subtype through wild bird populations [42–44]. Serology has proved particularly valuable in understanding HPAI H5 exposure in wild birds, notably in *Anas* ducks which have subclinical infections. For example, a recent study demonstrated repeated waves of HPAI H5N1 in waterfowl in eastern Canada [45]. Critically, these waves were revealed only through serological monitoring; as ducks were not dying they were not sampled via passive surveillance [45]. Indeed, it is only through serology that the true extent of viral spread, infection burden, and survival of HPAI H5N1 can be evaluated [*e.g.* [46–48]. Finally, serology can further provide a critical component of surveillance systems in situations where it is challenging to undertake active surveillance at high temporal resolution as required when targeting active infections and estimating viral prevalence.

However, key challenges to integrating serology remain. For example, the immune responses in many wildlife species are not well described, such that beyond “poor immune memory” [49], we don’t have a good understanding of antibody longevity in wild birds. From a study in geese, anti-NP antibodies were detectable for approximately one year [50]. Studies of repeatedly sampled Mallards indicated sero-reversion of anti-NP antibodies approximately 6 months after their final infection [51]. Anti-HA antibodies appear to wane much faster, with a measurable decrease in anti-HA antibodies by 45 days post vaccination in Mallards [52]. As such, it is unlikely that the between-year differences we describe in this study are due to antibody persistence over years, but are rather reflecting infection load in the previous 2-12 months. Thus, for serological data to better inform reconstructing LPAI epidemiology, more detailed studies on the longevity of the various antibodies in key LPAI-reservoir species would be required.

Overall, the emergence of HPAI H7 is driven by a complex interplay of environmental (e.g. water in landscape), host-specific (e.g. duck population dynamics), virological (e.g. LPAI diversity), and anthropogenic (e.g. characteristics of poultry operations) factors. While the presence of LPAI H7 in wild bird populations provides a continuous source of potential outbreaks, the chance of the virus spilling over into poultry and evolving into a high pathogenicity strain is promoted under specific conditions only. Key triggers include environmental factors such as rainfall and drought cycles that influence wild bird behaviour and viral transmission dynamics, as well as genetic mutations or reassortment events that increase virulence potential. Spillover from wild birds to poultry can occur when these factors converge. The unpredictability of influenza virus evolution, combined with limited comprehensive real-time surveillance, makes it difficult to pinpoint precisely when spillover may occur and why HPAI H7 emerges when it does. Understanding these dynamics in more detail to allow better risk mitigation, would require ongoing fine-scale surveillance, both in wild birds and poultry, and better ecological models to predict and prevent future virus spillover and subsequent outbreaks.

## Acknowledgements

We would like to thank the many individuals and organisations that contributed to the capture and sampling of ducks, and would notably extend our gratitude towards the Game Management Authority, Victorian Wader Study Group, Trent Leen and Toby Ross. We thank the jurisdictions, programs, and individuals responsible for generating and sharing qPCR and sequencing data under the NAIWB program. We would like to acknowledge ACDP for sharing H7 antiserum, Yi-Mo Deng and Hilda Lau for sequencing support, and Wytamma Wirth for assistance with Bayesian skygrid modelling. We would like to acknowledge Ivano Broz for depositing sequences in GenBank.

## Ethics statement

Ethics approval is not required for faecal environmental and hunter shot duck samples collected by state surveillance (Victoria, NSW, South Australia, ACT). For samples from live captured ducks, research was conducted under approval of Deakin University Animal Ethics Committee (application numbers B39-2019 and B28-2023), Wildlife Act 1975 Research Authorisation from the Victorian Department of Energy, Environment and Climate Action (permits 10009534, 10010870, and 10011005) and the Australian Bird and Bat Banding Scheme (banding authority 2915).

## Data Availability

All H7 sequences requested from the NAIWB program, as well as the H7 genome generated in this study, have been deposited in GenBank (Accessions: PX124824-PX124912). Underlying data have been made available at (https://github.com/michellewille2/H7Manuscript)

